# Super-resolved local recruitment of CLDN5 to filtration slits implicates a direct relationship with podocyte foot process effacement

**DOI:** 10.1101/2021.02.02.429338

**Authors:** Florian Tesch, Florian Siegerist, Eleonora Hay, Nadine Artelt, Christoph Daniel, Kerstin Amann, Uwe Zimmermann, Panagiotis Kavvadas, Olaf Grisk, Christos Chadjichristos, Karlhans Endlich, Christos Chatziantoniou, Nicole Endlich

## Abstract

Under healthy conditions, foot processes of neighboring podocytes are interdigitating and connected by an electron-dense slit diaphragm. Beside slit diaphragm proteins, typical adherens junction proteins are also found to be expressed at this cell-cell junction. It is therefore considered as a highly specialized type of adherens junction.

During podocyte injury, podocyte foot processes lose their characteristic 3D structure and the filtration slits typical meandering structure gets linearized. It is still under debate how this change of structure leads to the phenomenon of proteinuria. Using super-resolution 3D structured illumination microscopy, we observed a spatially restricted up-regulation of the tight junction protein claudin 5 (CLDN5) in areas where podocyte processes of patients suffering from minimal change disease (MCD), focal and segmental glomerulosclerosis (FSGS) as well as in murine nephrotoxic serum (NTS) nephritis and uninephrectomy DOCA-salt hypertension models, were locally injured. CLDN5/nephrin ratios in human glomerulopathies and NTS-treated mice were significantly higher compared to controls. In patients, the CLDN5/nephrin ratio is significantly correlated with the filtration slit density as a foot process effacement marker, confirming a direct association of local CLDN5 up-regulation in injured foot processes. Moreover, CLDN5 up-regulation was observed in some areas of high filtration slit density, suggesting that CLND5 up-regulation preceded the changes of foot processes. Therefore, CLDN5 could serve as a biomarker predicting early foot process effacement.

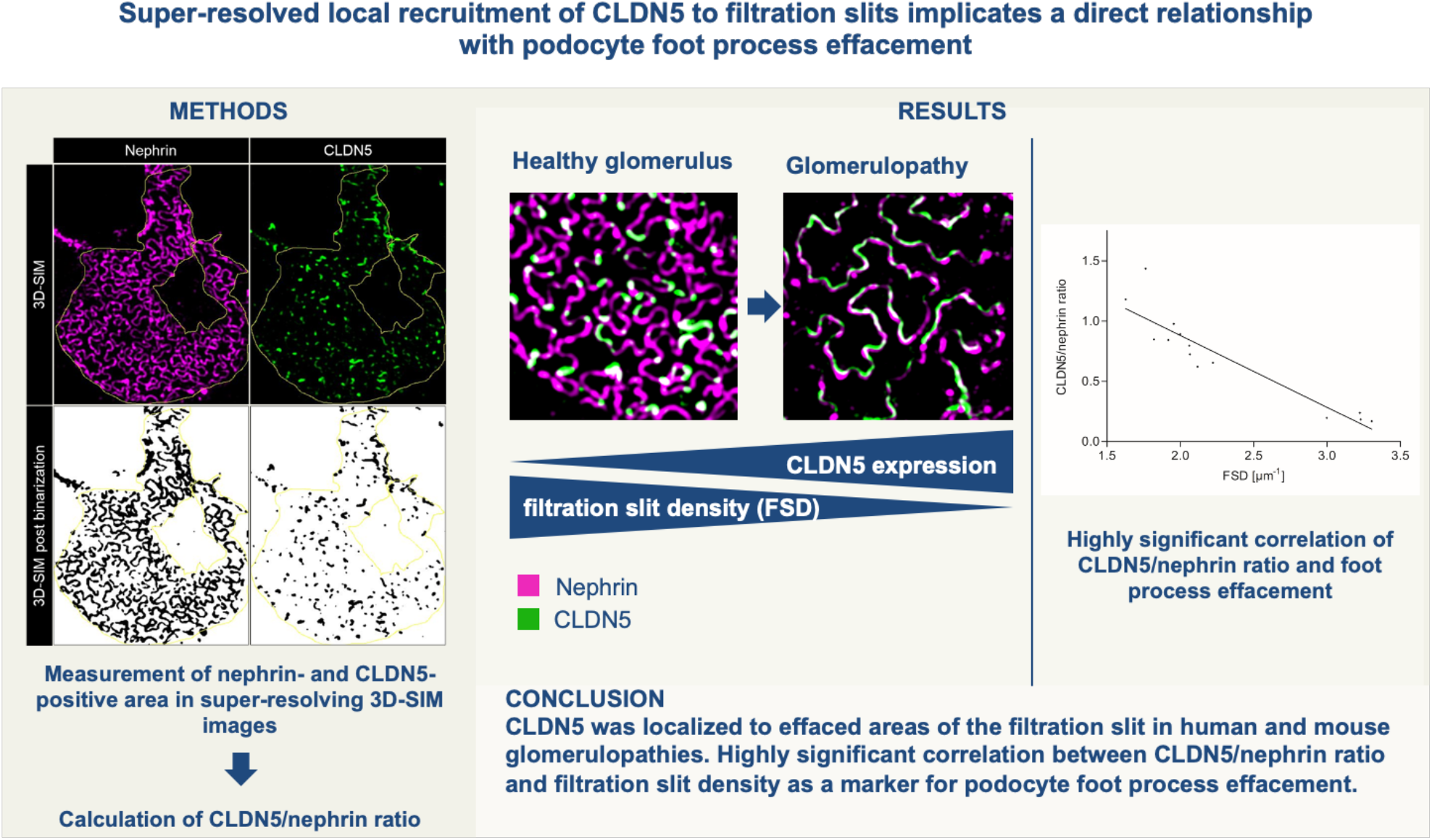

## Introduction

Interdigitating podocyte foot processes are an essential part of the kidney filtration barrier. The filtration slit (FS) between foot processes is bridged by the slit diaphragm (SD) which is structured by homodimerization of the transmembrane protein nephrin (8). At early time points during glomerular development, podocyte precursor cells are connected by tight junctions (13). Later, at the capillary loop stage, the number of occluding junctions reduces, and the SD is finally formed. Additionally, other filtration slit associated proteins like NEPH1 (4), podocin (1), CD2AP (15) or mFAT1 (3), which are essential for the size selectivity of the glomerular filtration barrier, are expressed in podocytes and localize to the SD.

This unique cell-cell contact was also referred by Farquahr et al. as a specialized type of adherens junction (2). Farquhar and coworkers have further shown that after the onset of a glomerular disease, which is associated with foot processes effacement, the composition of this specific cell-cell contact changes and converts back to occluding junctions indicating a de-differentiation of this postmitotic cell type (2, 13). These morphological changes of the foot processes could be studied in detail by electron microscopy in the past, however the exact molecular composition of these occluding junctions in interdigitating podocytes remained unclear for decades. In 2009, it was demonstrated that the tight junction proteins JAM-A, occludin and cingulin are expressed at the FS of healthy rat podocytes (5). Additionally, in puromycin aminonucleoside nephropathy (PAN) rats, a well-established model to induce podocyte foot process effacement, these tight junction proteins were significantly upregulated. This suggested a basic mechanism of tight junction formation in nephrotic kidneys.

Beside these results, it was reported that claudin 5 (CLDN5), a tetraspanning transmembrane protein of the claudin family (6, 17), is a major component of tight junction complexes. Koda and coworkers who investigated CLDN5 in rats and mice (9) have clearly shown by *in situ* hybridization that CLDN5 mRNA is expressed in podocytes, in endothelial cells of the arterioles at the glomerular vascular pole as well as in the endothelium of larger arteries. Transmission electron microscopy (TEM) images further showed that CLDN5 is localized at the SD of control rats as well as of rats with PAN-induced foot process effacement. Surprisingly, Western blot analysis of isolated glomeruli revealed that CLDN5 is not upregulated after PAN treatment. This is contrastive to other SD-associated tight junction proteins like JAM-A, occludin and cingulin that were significantly regulated in glomerular disease (5).

In this present study we wanted to analyze whether we can resolve the localization of a podocyte-specific claudin on the level of individual foot processes. Furthermore, we wanted to prove our hypothesis that the *de novo* localization of the tight junction protein CLDN5 precedes structural changes of podocyte foot processes and therefore indicates the dedifferentiation of podocytes.

Since healthy podocyte foot processes are at the borderline of resolvability of conventional (fluorescence) light microscopes, which is limited to 200 nm in the lateral direction, electron microscopy has been the gold standard for its visualization for decades. However, this technique is time consuming, highly sophisticated and can only analyze a thin layer of 50-90 nm of a kidney section. To overcome this obstacle, our group has established the 3D analysis of podocyte FS by the super-resolution microscopy technique 3D-SIM with 4 μm paraffin embedded tissue sections obtained from the diagnostic routine (16).

Herein, we revealed by 3D-SIM that the podocyte-specific CLDN5 has a specific staining pattern in healthy and diseased kidneys that can be resolved by 3D-SIM, Moreover, we suggest CLDN5 as a possible early effacement marker.

## Results

### CLDN5 is expressed in podocyte foot processes

To investigate which claudins are expressed in a podocyte-specific manner, we performed a database analysis using the Kidney Cell Explorer (https://cello.shinyapps.io/kidneycellexplorer/) created by Ransick and colleagues which is based on a single-cell RNA sequencing dataset of murine kidneys (12). Of all claudins annotated in the database, we found that Claudin-5 (CLDN5) transcripts specifically clustered in the podocyte population whereas the other claudins were expressed more distally in the nephron (Fig. 1). We verified this finding in a microarray dataset (http://www.nephroseq.org 02/2020, University of Michigan, Ann Arbor, MI) of human isolated glomeruli, in which CLDN5 was statistically significant enriched in glomeruli in comparison to the tubular kidney fraction (10).

**Figure 1:**
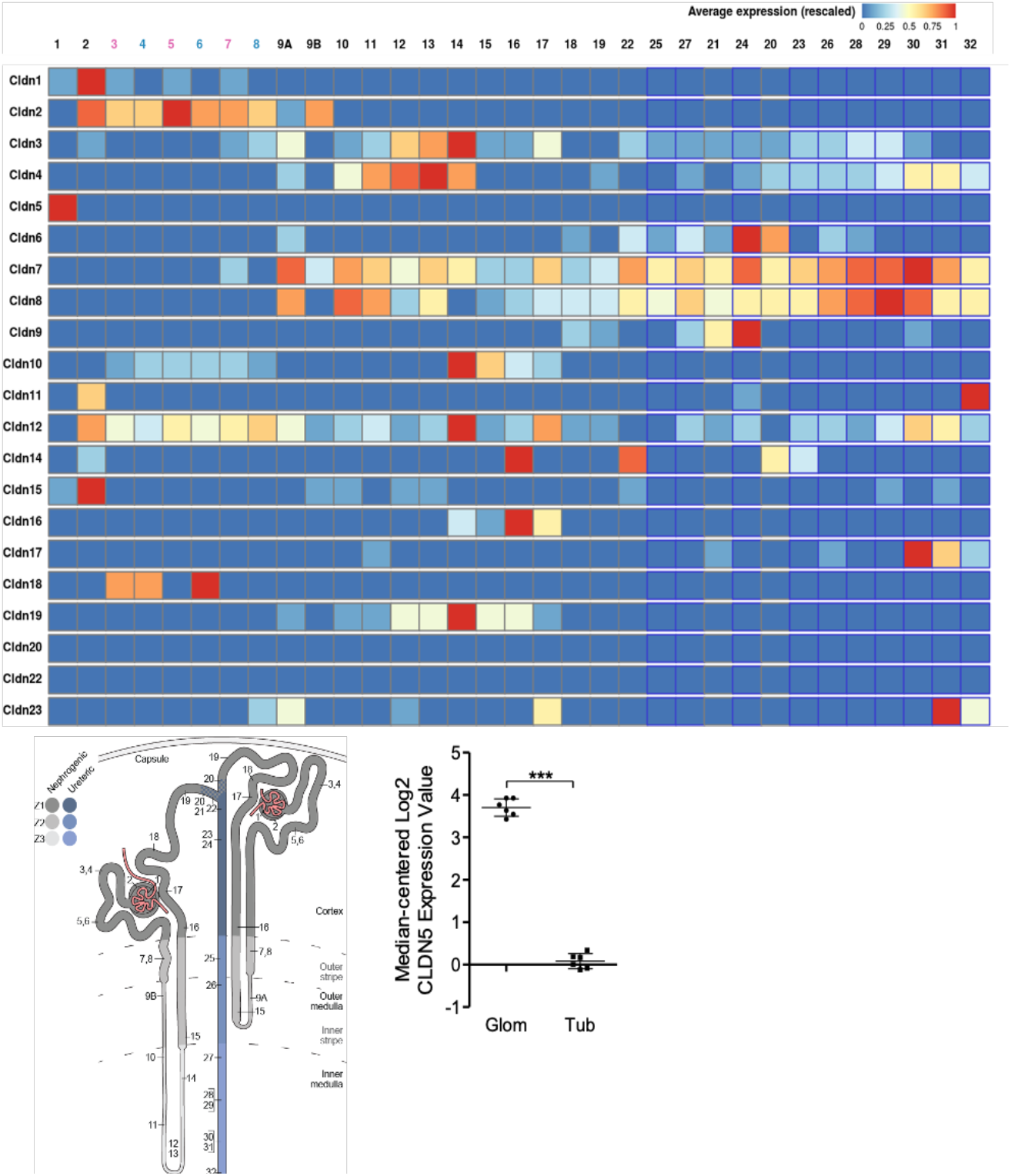
Database analysis of CLDN1 to CLDN23 using the Kidney Cell Explorer by Ransick and colleagues (12) based on a single-cell RNA sequencing dataset of murine kidneys to determine podocyte-specific claudins. CLDN5 is strongly expressed in ID 1 of the nephron representing podocytes. Other claudins were expressed further distally in the nephron or not expressed at all. The results were verified by analysis of the CLDN5 expression based on isolated glomeruli by Lindenmeyer et al. (10) which showed a significantly higher median-centered Log2 CLDN5 expression in the glomerulus compared to the tubule apparatus. *** P<0.001

Furthermore, the Proteinatlas database (18) (http://www.proteinatlas.org) was used and the expression of the five most proximal expressed claudins CLDN1, 2, 5, 11, 12 was evaluated by immunohistochemistry of kidney tissue (Fig. 2). As described before, we found that CLDN1 was expressed within parietal epithelial cells in contrast to CLDN5 which is expressed by cells on the glomerular tuft. As shown in Figure 2, anti-CLDN5 antibodies showed a linear staining pattern along the glomerular capillaries indicating a localization of CLDN5 in podocyte foot processes or at the slit diaphragm. The other claudins were expressed mainly non-glomerular and in the tubular system, respectively.

**Figure 2:**
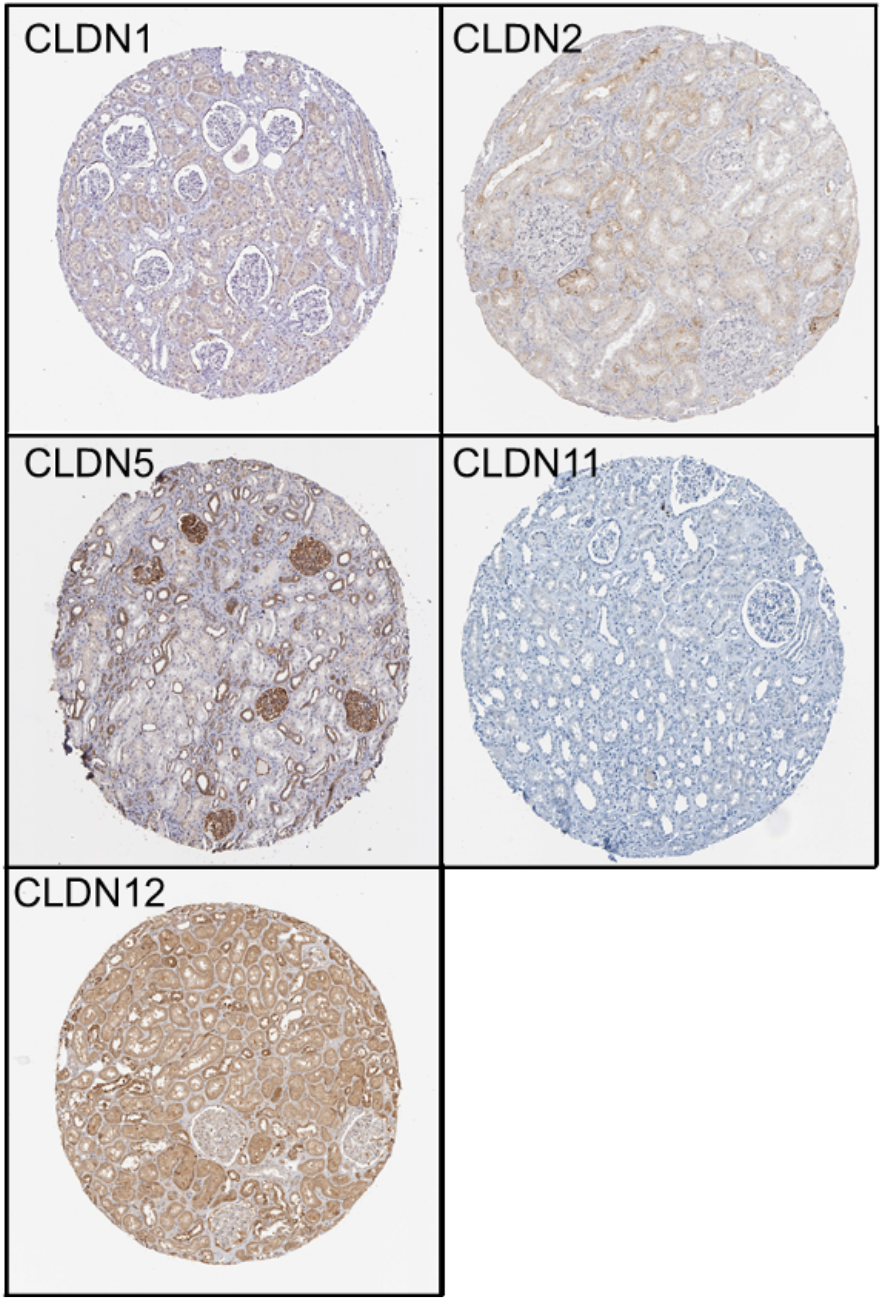
Evaluation of immunohistochemistry stainings of CLDN1, 2, 5, 11 and 12 from the Proteinatlas database (18) to investigate murine glomerular claudin expression. CLDN1 was expressed in parietal epithelial cells but not in podocytes. CLDN5 showed linear staining along the glomerular capillaries indicating localization to podocyte foot processes or the slit diaphragm. CLDN 2 and 12 were expressed mainly in the tubular system. CLDN 11 showed overall low antibody enhancement in kidney sections.

To further localize CLDN5 within the healthy kidney, we stained kidney sections of human nephrectomy samples and healthy wildtype mice with a monoclonal antibody directed against CLDN5 and detected this antibody with fluorescence-labeled secondary antibodies. In confocal laser scanning micrographs (C-LSM) of healthy human samples, we found CLDN5 localized at the podocyte slit membrane partially colocalized with the slit diaphragm protein nephrin, as well as the glomerular endothelium identified by a co-staining against CD31. Additionally, we found CLDN5 localized in the arterioles of the glomerular vascular pole. In sections of control mice, the CLDN5 expression was also found at the SD and vascular pole, but no expression was found in glomerular capillaries (Fig. 3).

**Figure 3.**
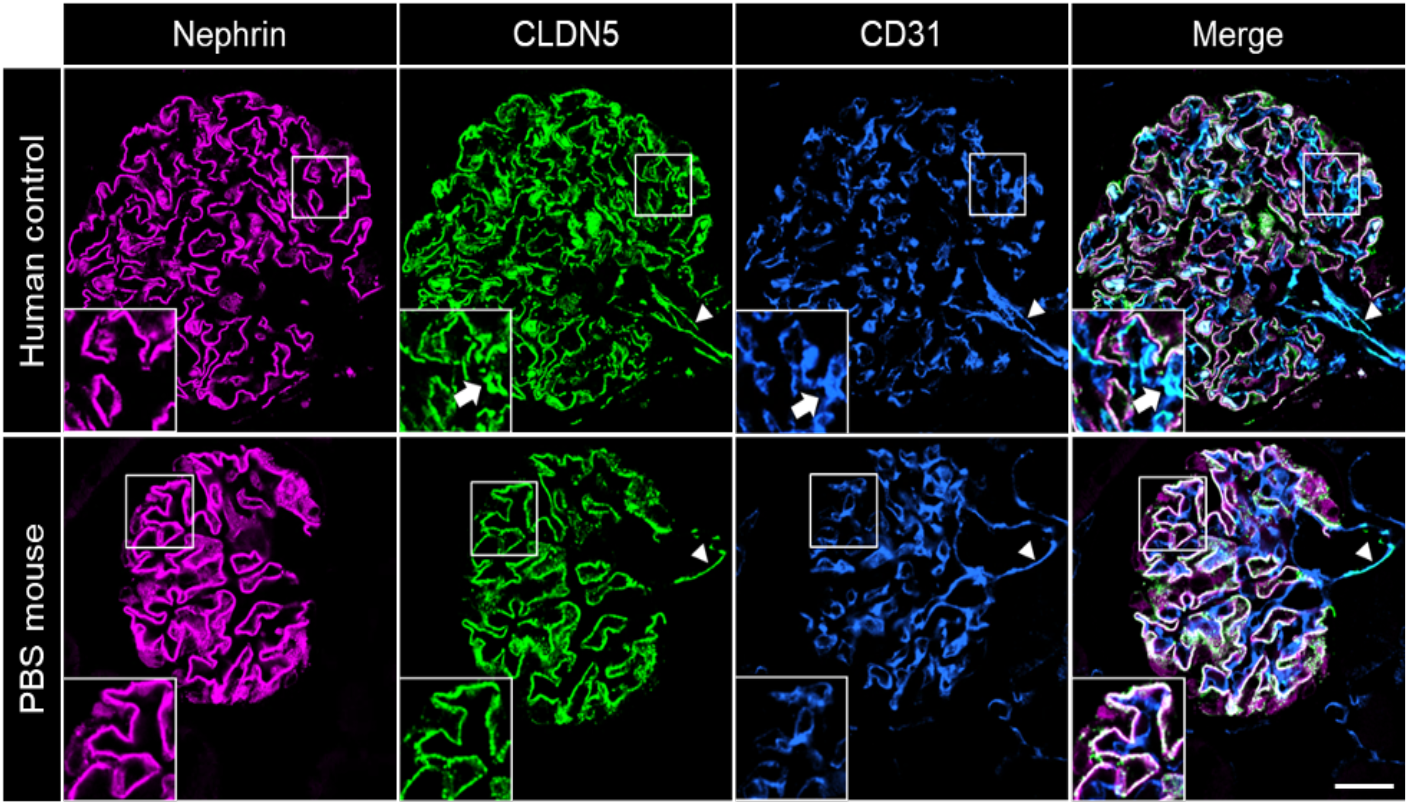
C-LSM images of healthy human glomeruli showed CLDN5 expression in the FS and the endothelium of glomerular capillaries (arrows) and of arterioles at the vascular pole (arrowheads). In mouse glomeruli CLDN5 was expressed in the FS and at the vascular pole (arro wheads), but no expression was found in glomerular capillaries. The scale bar indicates 20

### 3D-SIM studies of CLDN5

To further determine the subcellular localization of CLDN5 beyond the diffraction limit in podocytes, we used 3D-SIM of formalin fixed paraffin embedded (FFPE) tissue samples which allows high resolution fluorescence imaging of podocyte foot processes. With our experimental setup and sample preparation, we reached an xy-resolution down to ~115 nm which is sufficient to resolve individual healthy podocyte foot processes and the slit diaphragm (Fig. 4). In 3D-SIM images of healthy human and mouse samples CLDN5 focally co-localized with the slit diaphragm protein nephrin along the FS (Fig. 5, Fig. 6).

**Figure 4.**
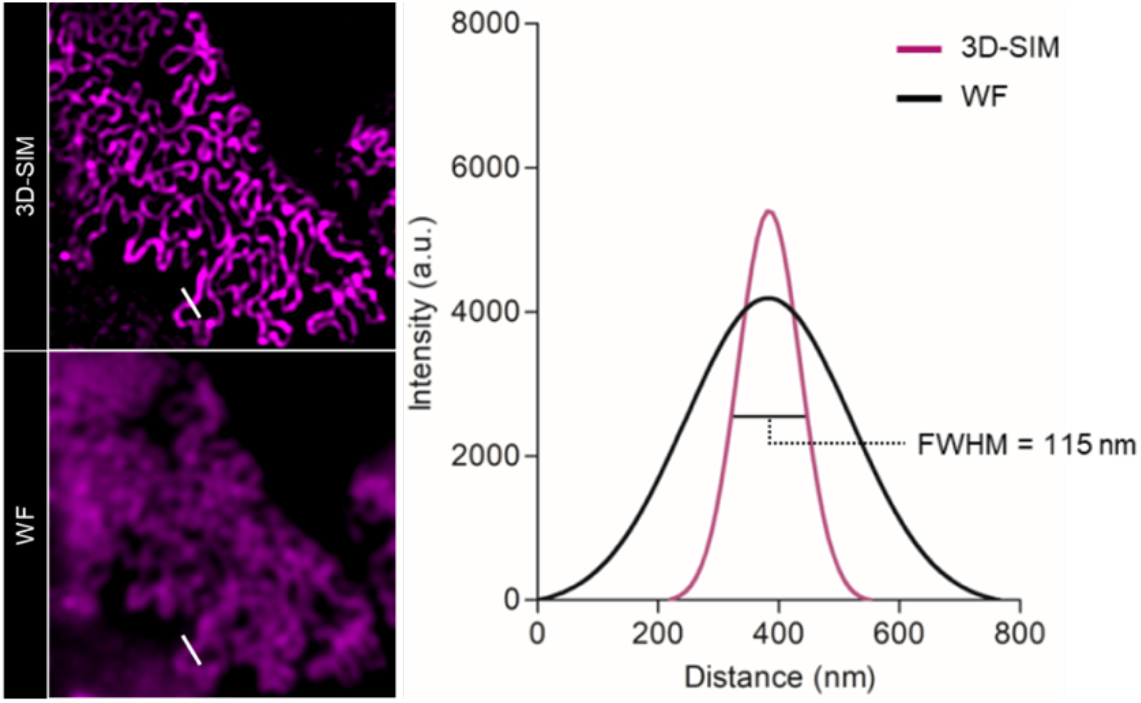
Micrographs of nephrin-stained human glomeruli showed increased resolution of 3D-SIM images compared to wide field (WF) images of the same z-stack. 3D-SIM enabled FS imaging with a xy-resolution of 115 nm (FWHM = full width at half maximum).

**Figure 5.**
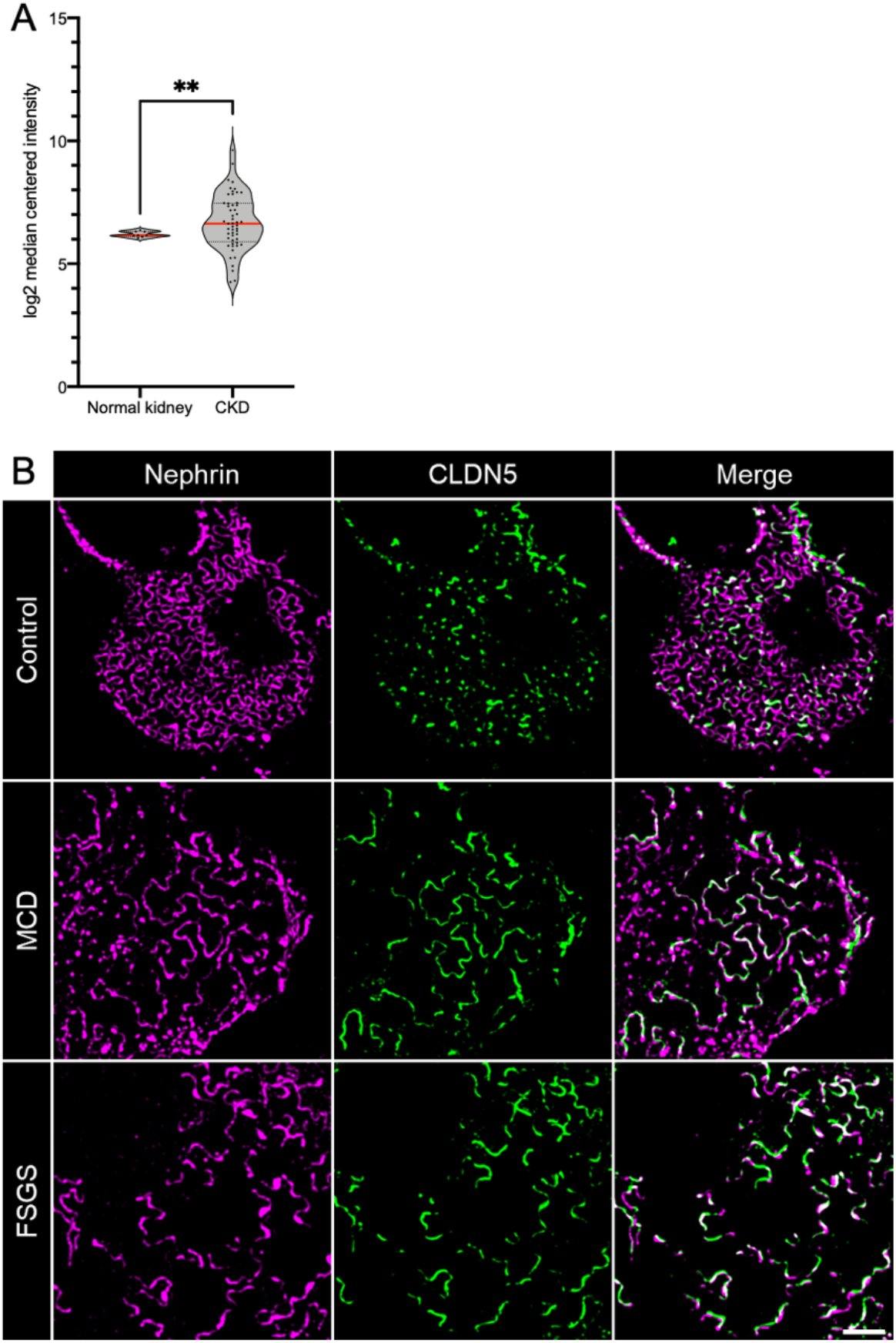
**(A)** CLDN5 expression analysis of CKD patients from the Nakagawa CKD microarray cohort (11). In CKD, CLDN5 was significantly upregulated 2.05-fold (P=0.005) compared to healthy donor kidneys. Panel **(B)** shows 3D-SIM images of human glomeruli. CLDN5 focally co-localized with the slit diaphragm protein nephrin along the FS in human nephrectomy samples. MCD and FSGS glomeruli showed extended CLDN5-positive lines at the FS in areas of effaced podocyte foot processes. The scale bar indicates 2 μm.

**Figure 6.**
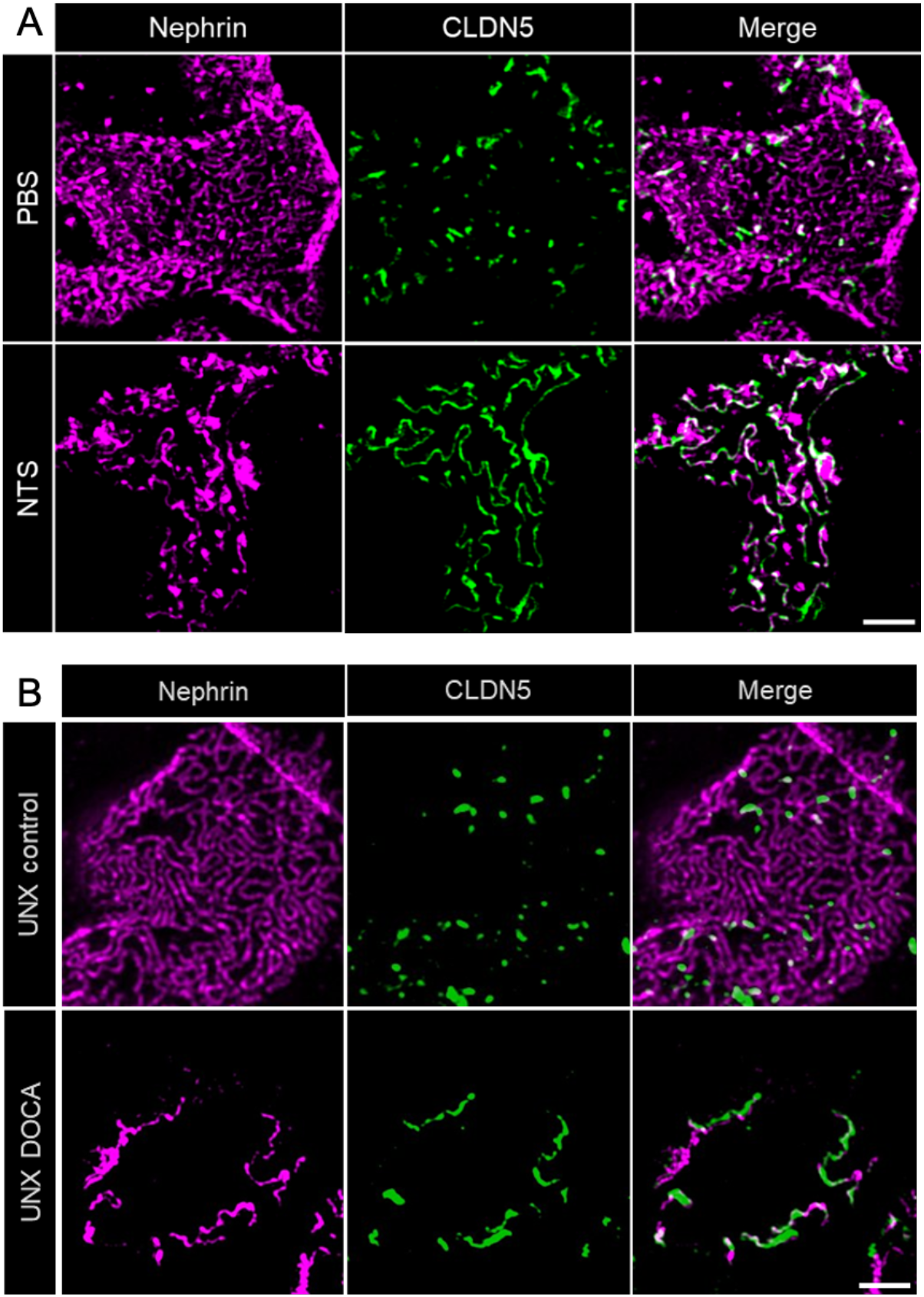
3D-SIM images of mouse glomeruli. **(A)** CLDN5 focally co-localized with nephrin along the FS in PBS-treated control mice. Similar to human biopsies, glomeruli of mice treated with nephrotoxic serum (NTS) showed extended CLDN5-positive lines at the FS in areas of effaced podocyte foot processes. Panel **(B)** shows focal CLDN5 localization to effaced FS in mice of the uninephrectomy deoxycorticosterone acetate (DOCA) salt hypertension mouse model compared to UNX controls. The scale bar indicates 2 μm.

### CLDN5 is upregulated in MCD and FSGS

To find out whether CLDN5 was regulated in glomerular disease, we screened disease-specific datasets within the aforementioned Nephroseq database. We found CLDN5 to be statistically significant upregulated 2.05-fold (P=0.005) in the Nakagawa CKD microarray cohort (11) compared to healthy donor kidneys (Fig. 5A).

Then, we co-immunostained biopsy specimen of MCD and FSGS patients for CLDN5 and nephrin. In contrast to healthy samples, patient biopsies showed extended CLDN5-positive lines that were localized in areas of effaced podocyte foot processes. Figure 5B shows more linearized and less meandering nephrin-positive filtrations slits which can be used as a direct marker for broadened foot processes.

### CLDN5 and nephrin are inversely expressed in murine models of glomerular disease

To investigate whether this upregulation of CLDN5 is a species-independent and general mechanism rather than a human disease specific phenomenon, we imaged FFPE sections of mice (10 weeks old) with induced nephrotoxic serum (NTS) nephritis 12 days after NTS-injection and corresponding control-treated mice. It is well established that after NTS-injection, mice develop severe glomerulonephritis with heavy proteinuria accompanied by severe podocyte foot process effacement. Similar to the human biopsies, a massive local upregulation of CLDN5 was found in the cell-cell junctions of effaced foot processes in NTS-injected mice (Fig. 6A).

Furthermore, to investigate if the upregulation of CLDN5 together with increasing foot process effacement, is a general rather than a model-specific phenomenon, we additionally evaluated sections of the uninephrectomy deoxycorticosterone acetate (DOCA)-salt hypertension mouse model. In this model, mice develop hypertension after uninephrectomy, parenteral DOCA application and enteral salt application. During the course of disease, mice develop progressive albuminuria and glomerulosclerosis similar to human secondary FSGS. As indicated in Figure 6B, we found that also in this model, CLDN5 was recruited to effaced FS. The local phenotype in effaced areas was similar to the previous findings while in general the phenotype in the DOCA-salt hypertension model was less uniform and more focally restricted to single glomeruli and not as distributed as in the NTS model.

Independent of the model or disease investigated, we observed that during the beginning of foot process effacement (phase 1), nephrin is still continuously localized at the FS followed by an upregulation of CLDN5. Interestingly, effacement without CLDN5 up-regulation could only be found very rarely in some FS areas. With progressing effacement (phase 2), nephrin staining appeared discontinuous. Additionally, CLDN5-positive but nephrin-negative FS areas were identified, especially in NTS-treated mice indicating the severity of the NTS-model with full dedifferentiation of the FS to a tight junction phenotype (Fig. 7).

**Figure 7.**
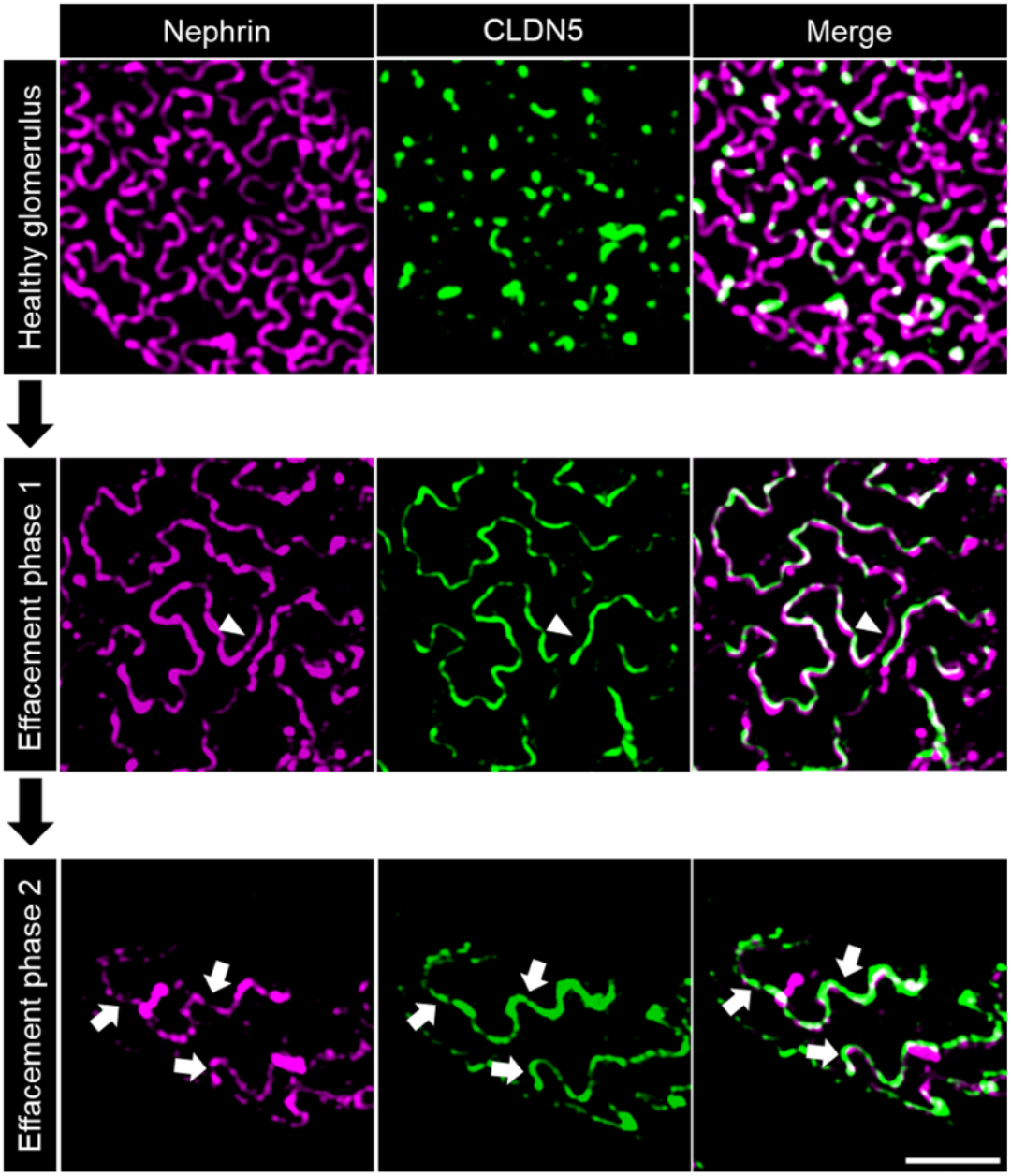
Evolution of CLDN5 upregulation in 3D-SIM images. Compared to healthy glomeruli, in phase 1 of effacement CLDN5 was upregulated following the nephrin-positive FS. Nephrin is continuous and effaced areas without CLDN5 could be observed (arrowheads). With progressing effacement (phase 2), nephrin became discontinuous and CLDN5-positive but nephrin-negative areas were found (arrows). The scale bar indicates 2 μm.

### CLDN5/nephrin ratio correlates with the FSD

These results raised the question, whether a CLDN5 up-regulation could only be observed in strongly effaced FS areas or if it could be also observed in areas with a high filtration slit density (FSD). While most healthy FS areas only showed focally localized CLDN5 expression, as described above, several FS areas with a high FSD (indicating healthy foot processes) seem to upregulate CLDN5 (Fig. 8).

**Figure 8.**
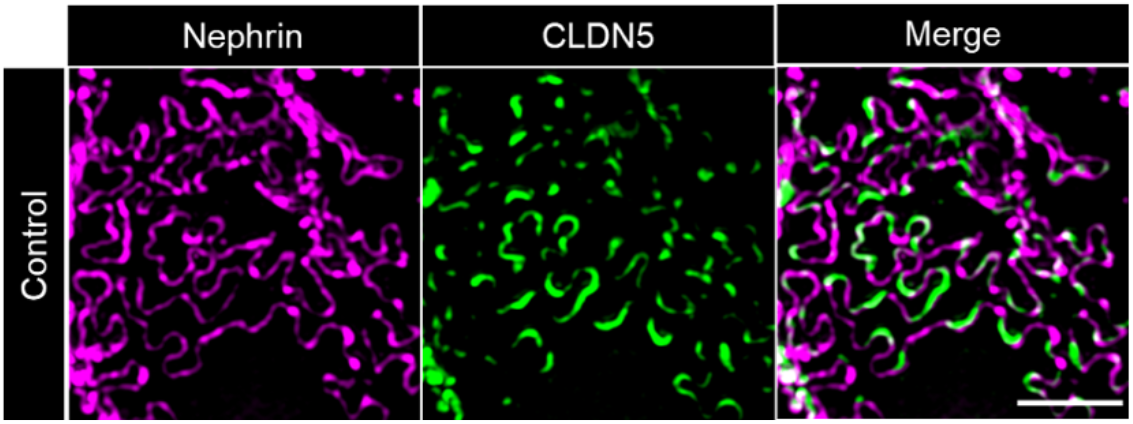
CLDN5 upregulation in non-effaced FS areas of human control glomeruli. In several FS areas CLDN5 forms continuous lines despite lack of foot process effacement and normal FSD. The scale bar indicates 2 μm.

To quantify these impressions, we developed a FIJI (14)-based macro to determine the CDLN5/nephrin ratio (Fig. 9). Z-stacks were processed into maximum intensity projections. After selection of a plan FS area, thresholding and binarization of the CLDN5- and nephrin-channel, we quantified the CLDN5- and nephrin-positive areas. This allowed calculation of the CLDN5/nephrin ratio.

**Figure 9.**
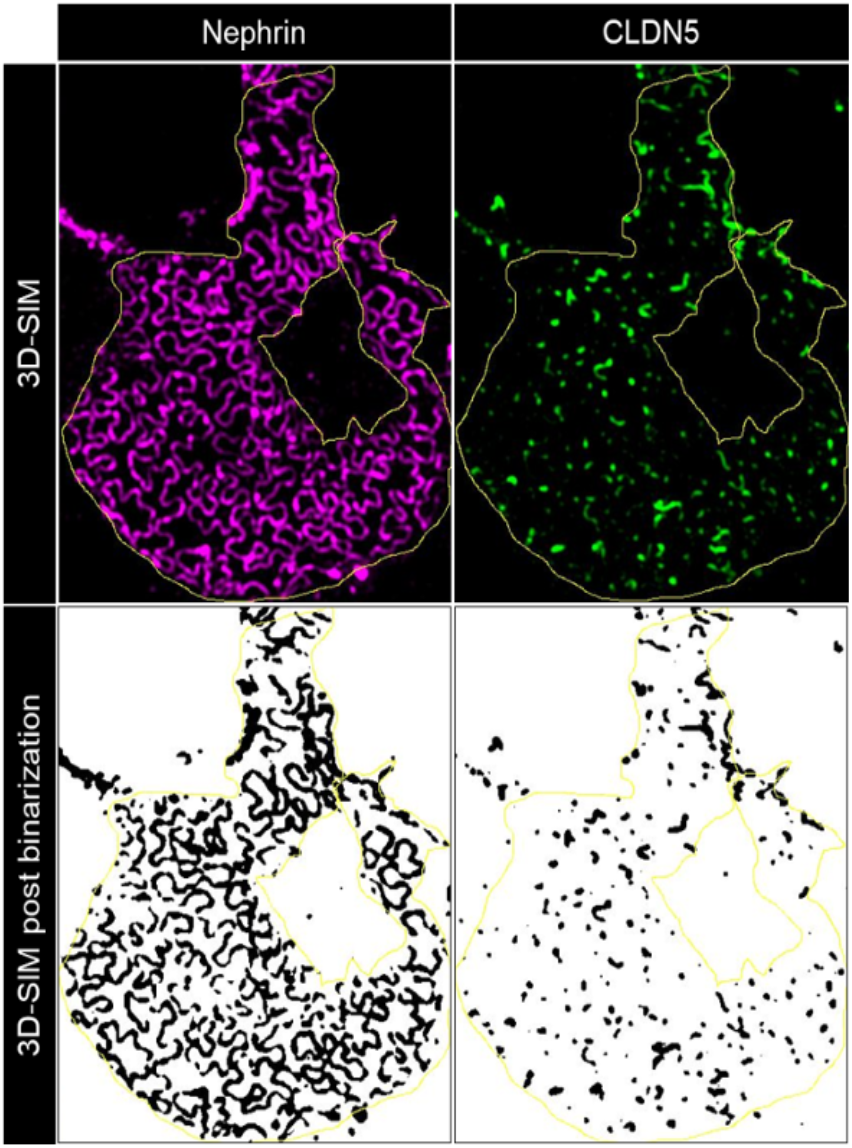
CLDN5/nephrin ratio measurement in maximum intensity projections of 3D-SIM images of a selected FS area. The nephrin- and CLDN5-channel were transformed into black-and-white images by binarization following defined thresholds.

As expected, we found a significantly higher CLDN5/nephrin ratio in MCD and FSGS samples than in human controls. These findings could be verified in mice as well: the CLDN5/nephrin ratio in NTS-injected mice was statistically significantly higher compared to PBS-injected control mice (Fig. 10).

**Figure 10.**
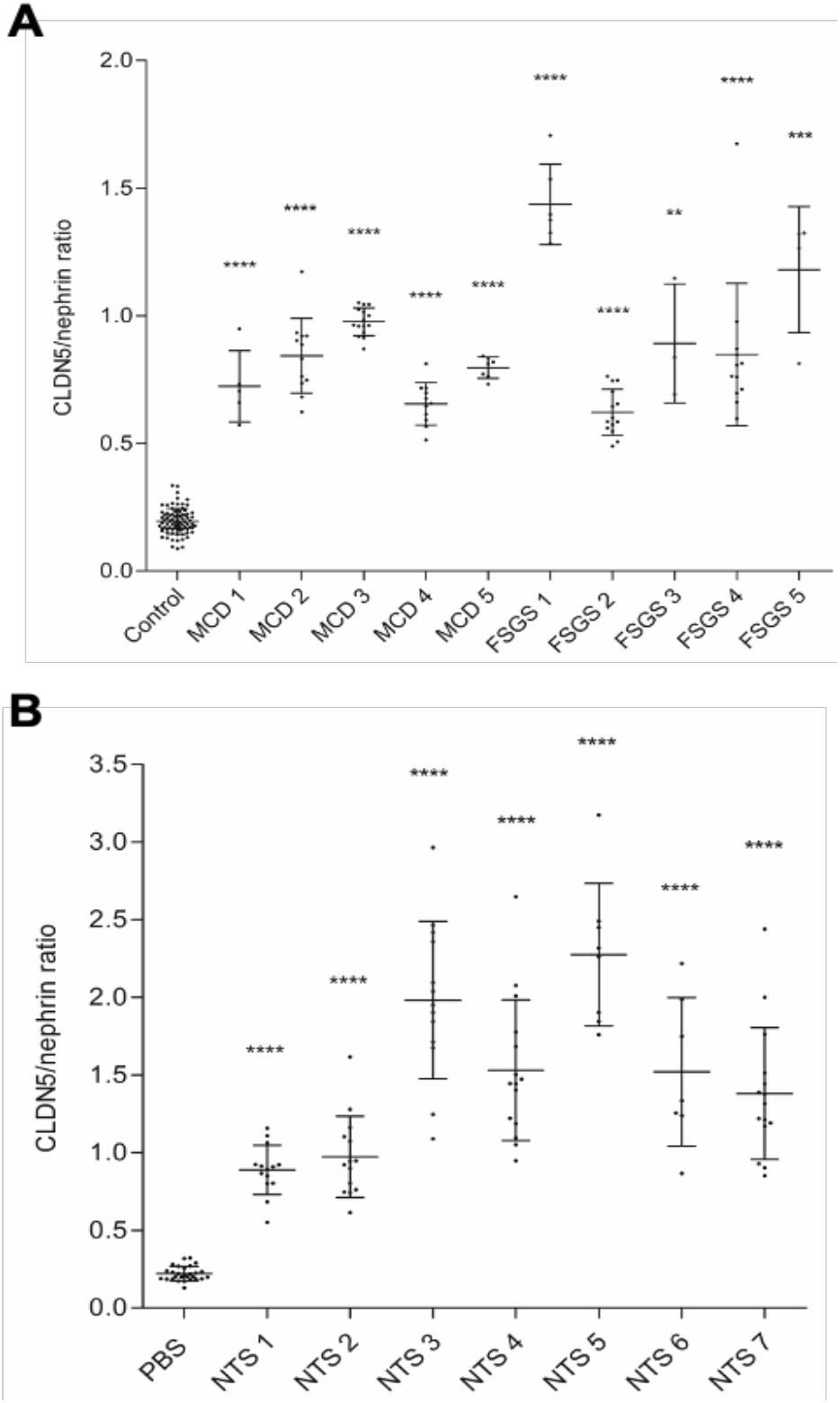
**(A)** Results of CLDN5/nephrin ratio measurement showed a significantly higher CLDN5/nephrin ratio in MCD (n=5) and FSGS (n=5) biopsies compared to human nephrectomy samples as controls (n=4). Panel **(B)** shows a significantly higher CLDN5/nephrin ratio in NTS-injected mice (n=7) compared to PBS-injected control mice (n=3). In **(A)** and **(B)** every measuring point represents the glomerular mean of at least 20 measured areas. Error bars show standard deviation. **** P<0.0001, *** P<0.001, ** P<0.01, Mann-Whitney *U* test.

The results showed a species-independent association of podocyte foot process effacement and CLDN5 up-regulation. To further investigate this association in detail, we measured the FSD with our established *Podocyte Exact Morphology Measurement Procedure* (PEMP) (16). PEMP allows a quick and exact quantification of the FSD of each glomerulus where the CLDN5/nephrin ratio as determined before. Thereby, PEMP quantified the total length of the nephrin-positive FS (I_FS_) in a selected FS area as well as the size of the area (A). The FSD was calculated by dividing I_FS_ by A.

By using PEMP, we found that the FSD in MCD and FSGS samples was statistically significant reduced compared to healthy human nephrectomy samples (Fig. 11).

**Figure 11.**
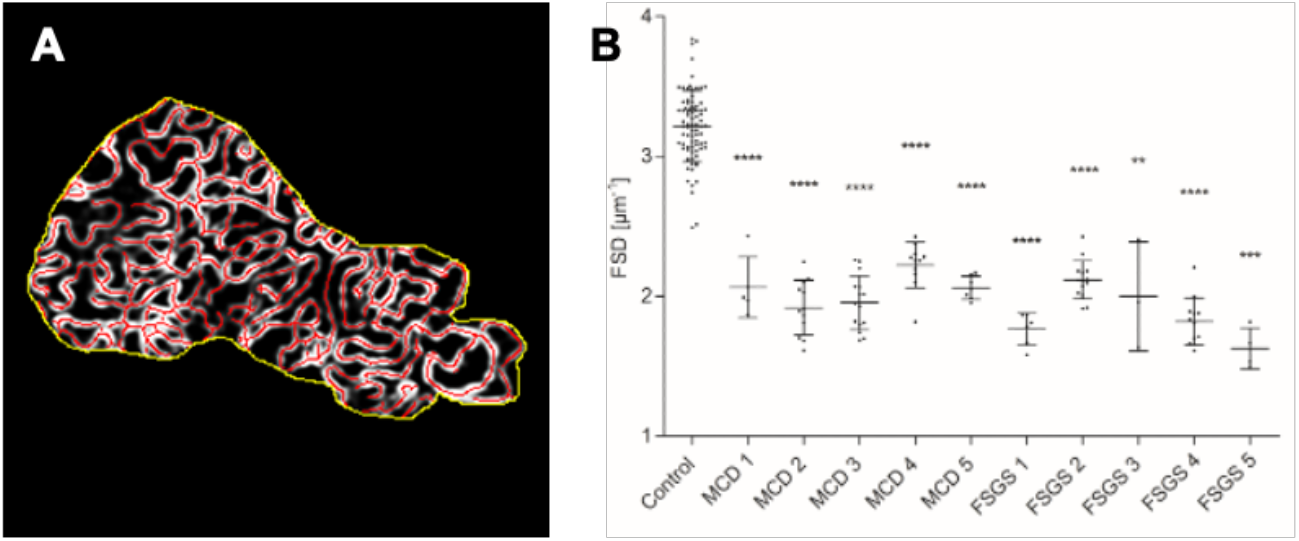
Measurement of the filtration slit density (FSD) in MCD (n=5) and FSGS (n=5) compared to human nephrectomy samples as controls (n=4) with PEMP. **(A)** shows automated detection of the nephrin-positive FS in a selected FS area. **(B)** MCD and FSGS biopsies showed a significantly reduced FSD compared to controls. Every measuring point represents the glomerular mean of at least 20 measured areas. Error bars show standard deviation. **** P<0.0001, *** P<0.001, ** P<0.01, Mann-Whitney *U* test.

As shown in Figure 12, CLDN5/nephrin ratio and FSD correlated significantly (r=−0.952, P<0.0001, R^2^=0.906) in human samples, indicating that the CLDN5 localization in relation to the FS increased the broader the foot processes became.

**Figure 12.**
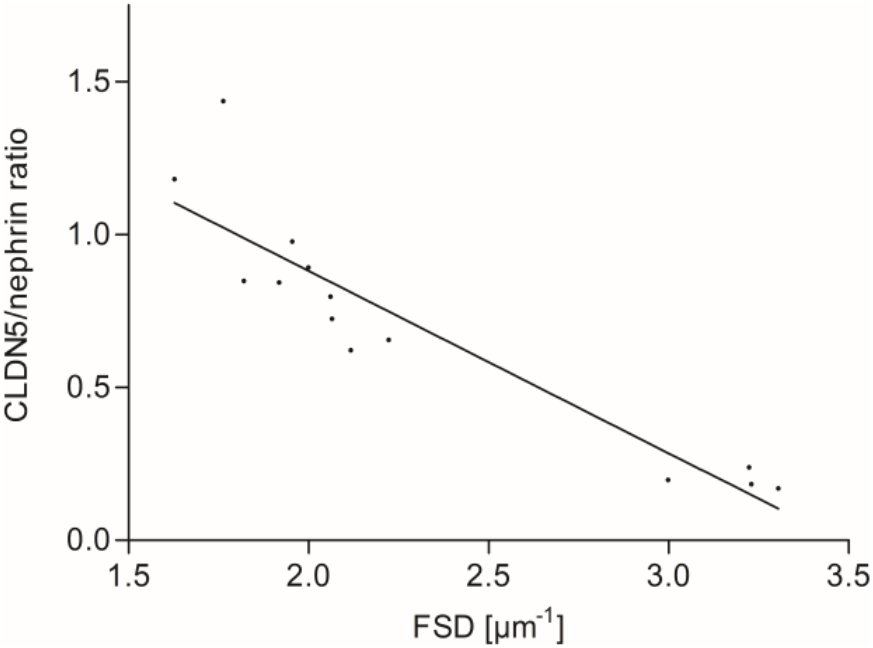
Correlation analysis of CLDN5/nephrin ratio and FSD in MCD (n=5), FSGS (n=5) and controls (n=4) showed a highly significant correlation of CLDN5/nephrin ratio and FSD. CLDN5 expression increased with increasing foot process effacement. Every measuring point represents the average for the whole sample. r=−0.952, P<0.0001, R^2^=0.906, Spearmans correlation.

## Discussion

To summarize, the super-resolving 3D-SIM technique revealed that CLDN5 was localized to those areas of the FS where foot processes were effaced. The CLDN5/nephrin ratio was significantly correlated with the FSD in human sections. We performed a super-resolution microscopy study of CLDN5 expression in human and mouse for the first time.

In our findings in mice, CLDN5 was not expressed in the capillary endothelium, but in the endothelia of arterioles of the vascular pole which was previously observed in rats as well (9). We found CLDN5 up-regulation in NTS mice compared to controls while Western blot analysis of CLDN5 expression in PAN-treated rats showed no up-regulation compared to controls (9). This may be traced back to the strong CLDN5 expression in arterioles of the vascular pole. Slight CLDN5 upregulation in the FS of PAN-treated rats could not be detected by Western blot most probably because of superimposing CLDN5 expression in arterioles of isolated glomeruli.

This superimposing effect could be even stronger in human sections because of the additional CLDN5 expression in glomerular endothelial cells. In this context, one major advantage of our 3D-SIM approach combined with FIJI-based evaluation is that CLDN5-up-regulation in the FS can be studied separately from endothelial CLDN5 expression.

Furthermore, we confirmed the basic mechanism of tight junction formation in the FS in kidney injury for CLDN5. Tight junction protein up-regulation in the FS has been shown in PAN-treated rats before for occludin, cingulin and JAM-A (5). Lately, 3D-reconstruction of FIB/SEM images showed a co-localization of slit diaphragms and tight junctions in rats on day 4 after PAN treatment as well (7): as initial alteration slit diaphragms moved further to the luminal side of the FS while tight junctions formed below. In this stage, SDs and tight junctions coexisted in close proximity. On day 8 after PAN treatment, slit diaphragms were replaced by tight junctions which connected strongly effaced foot processes and slit diaphragms vanished. This could be an explanation for our finding that CLDN5 and nephrin were co-expressed, that the podocyte cell-cell contact shows tight junction formation with an up-regulation of CLDN5 while the nephrin-containing slit diaphragm still exists more luminal.

In our study of MCD and FSGS biopsies, continuous nephrin lines as well as a nephrin-CLDN5 co-localization could be seen despite effacement. It is conceivable that kidney injury in MCD and FSGS was not as strong as injury in rats following PAN treatment. In mice, NTS injection led to severe foot process effacement and fragmented nephrin lines while CLDN5 was continuous in these areas. For this reason, in several NTS mice CLDN5/nephrin ratios distinctly greater than 1 were measured indicating destruction of slit diaphragm and locally complete tight-junction transformation. By investigating CLDN5 expression in DOCA-treated unilateral nephrectomy mice, we could confirm CLDN5 recruitment to effaced FS areas as a general mechanism of murine kidney injury, rather than an NTS-specific phenomenon. Taken together, CLDN5 upregulation was observed species- and model-independently in effaced FS areas.

In addition, our data showed FS areas of high FSD, where up-regulation of CLDN5 was observed. Importantly, in these areas CLDN5 up-regulation was found without significant foot process effacement. While in general CLDN5 expression increased with increasing effacement, these findings show that CLDN5 formation at the FS can precede the morphologic process of foot process effacement. For this reason, CLDN5 could be described as a potential early effacement marker predicting changes in foot process morphology. These results could only be discovered in control sections, because the MCD and FSGS samples we used were already globally effaced.

As explained above, Ichimura and colleagues demonstrated a distinct vertical distance between slit diaphragm and tight junction in the FS. To eliminate the influence of CLDN5 and nephrin being localized in slightly different planes of the z-stack maximum intensity projections were used. Manual creation of maximum intensity projections guarantees that no CLDN5-signal from capillary endothelium is projected into the FS area, which would otherwise lead to falsely high CLDN5/nephrin ratios. To avoid selection bias, CLDN5/nephrin ratio and FSD were measured in all evaluable FS areas. Only glomeruli with a minimum of 20 evaluable FS areas were included in this study.

In conclusion, our study showed CLDN5 localization to effaced areas of the FS in human and mouse glomerulopathies. Highly significantly correlation between FSD and CLDN5/nephrin ratio in human sections could be demonstrated. Our results show that with increasing effacement CLDN5 expression in the FS is increased simultaneously, while in several FS areas CLDN5 up-regulation preceded foot process effacement. CLDN5 can therefore be described as a marker predicting early foot process effacement. Our results hereby provide a better understanding of the sequence of tight junction formation in kidney injury.

## Materials and methods

### Kidney tissue

For our study, formalin-fixed paraffin-embedded (FFPE) human kidney biopsies of patients diagnosed with MCD (n=5) and FSGS (n=5) by experienced pathologists of the Department of Nephropathology of the University Erlangen-Nuremberg were used. As healthy controls, anonymized excess normal kidney tissue of partial nephrectomies of the Department of Urology of the University Medicine Greifswald was used (n=4). The use of the biopsies from Erlangen has been approved by the Ethics Committee of the Friedrich Alexander University of Erlangen-Nuremberg, waiving the need for retrospective consent for the use of archived excess material (Ref. No. 4415). The use of this excess kidney tissue from Greifswald has been approved by the Ethics Committee of the University Medicine Greifswald (Ref. No. BB 075/14). All subjects stated written informed consent. All experiments were performed in accordance with local guidelines overseen by the University Medicine Greifswald and University Greifswald, Greifswald, Mecklenburg - Western Pomerania. 4 μm sections were cut and mounted on Superfrost slides (R. Langenbrinck GmbH, Emmendingen, Germany).

Ten-week-old male C57BL/6 mice were retro-orbitally injected with 17 μl of nephrotoxic serum (NTS) per gram of body weight over 2 consecutive days (n=7) or with PBS as control (n=3). On day 12, mice were anesthetized with ketamine (100 mg/kg body weight)-xylazine (10 mg/kg body weight), sacrificed and kidneys were harvested, formalin-fixed and paraffin-embedded. 4 μm sections were cut and mounted on #1.5 High Precision coverslips (Paul Marienfeld GmbH, Lauda-Königshofen, Germany). All animal experiments were performed in strict accordance with good animal practice and animal research guidelines and were approved by the appropriate committee of the Institut national de la santé et de la recherche médicale (INSERM) and the Sorbonne Université Paris, France (Ref. No. B752001). FFPE kidney tissue of the uninephrectomy DOCA salt hypertension mouse model of a previous study was used which was prepared as described before (Schordan et al. 2013).

### Immunofluorescence staining

After deparaffinization in xylene and rehydration in a descending ethanol series, all sections were boiled with a pressure cooker in Tris EDTA buffer (10 mM Tris, 1 mM EDTA, bring to pH=9 using NaOH, add 500 μl Tween 20) for antigen retrieval. After autofluorescence quenching in 100 mM glycine diluted in purified water and three times washing in PBS, slices were incubated for 1 hour in blocking solution (1% FBS, 1% BSA, 0.1% fish gelatin, 1% normal goat serum in PBS). The primary antibodies (1:400 in blocking solution, polyclonal guinea pig anti-nephrin IgG GP-N2, Progen, Heidelberg, Germany; 1:400 in blocking solution, monoclonal mouse anti-Claudin-5 IgG, 4C3C2, Thermo Fisher Scientific, Waltham, Massachusetts, USA; 1:50 in blocking solution, polyclonal rabbit anti-CD31 IgG, ab28364, Abcam, Cambridge, UK) were incubated at 4°C on the slides overnight. The next day after five times washing in PBS and blocking for 45 minutes the secondary antibodies (all 1:600 in blocking solution: Cy3-conjugated donkey anti-guinea pig IgG (H+L), 706-165-148, Jackson Immuno Research, Hamburg, Germany; Alexa Fluor 488-conjugated F(ab)2-fragment goat anti-mouse IgG, 115-546-072, Jackson Immuno Research, Hamburg, Germany; Alexa Fluor 647-conjugated F(ab)2-fragment goat anti-rabbit IgG, 111-607-047, Jackson Immuno Research, Hamburg, Germany) were incubated at 4°C for 45 minutes, followed by five times washing in PBS and transfer to purified water. The slices were mounted in Mowiol for Microscopy (Carl Roth, Karlsruhe, Germany) using High Precision cover glasses (Paul Marienfeld GmbH, Lauda-Königshofen, Germany).

### Microscopy

For confocal laser scanning microscopy, a Leica TCS SP5 (Leica Microsystems, Wetzlar, Germany) equipped with a 63x (NA 1.4) oil immersion objective was used. Mouse glomeruli were imaged with 6.080 pixel/μm. Human glomeruli were imaged with 5.853 pixel/μm. For 3D-SIM a Zeiss Elyra PS.1 System (Zeiss Microscopy, Jena, Germany) equipped with a 63x (NA 1.4) oil immersion objective was used. Z-Stacks with a size of 1280 μm x 1280 μm with a slice-to-slice distance of 0.2 μm were acquired over approximately 4 μm using the 561 nm laser, (3% laser power, exposure time:100 ms) and the 488 nm laser (4% laser power, exposure time: 100 ms). The 28 μm period grating was shifted 5 times and rotated 5 times on every frame whilst acquiring widefield images. The 3D-SIM reconstruction was performed with the Zeiss ZEN Black Software using following parameters: Baseline Cut, SR Frequency Weighting: 1.0; Noise Filter: −5.6; Sectioning: 96, 81, 81. ZEN Blue software was used for maximum intensity projections.

### CLDN5/nephrin ratio measurement

Z-stacks of glomeruli stained for nephrin and CLDN5 were transformed into maximum intensity projections. For automated measurement of the CLDN5/nephrin ratio a customized macro for the ImageJ-based platform FIJI was programmed. A FS area is selected manually. The macro performs binarization of the nephrin-Cy3- and the CLDN5-Alexa488-channel according to defined thresholds creating black-and-white images. The nephrin- and CLDN5-positive area is measured and saved as an Excel file. With this data the CLDN5/nephrin ratio of the area was calculated. The CLDN5/nephrin ratio was evaluated for a minimum of 20 areas per glomerulus and the mean was calculated. To check for statistical difference between effaced kidney samples and controls, not completely normally distributed values were compared using Mann-Whitney *U* test using SPSS Statistics (25.0 IBM SPSS Inc., Chicago, IL, USA).

### Correlation of CLDN5/nephrin ratio with FSD

Filtration slit density (FSD) was measured with Podocyte Effacement Measurement Procedure (PEMP) as published before (16). Correlation analysis between CLDN5/nephrin ratio and FSD was performed using Spearmans correlation for not normally distributed data using SPSS Statistics. All graphs were set up with Prism 5.01 (GraphPad, CA, USA).

## Author contributions

F.T. performed experiments, analyzed data, prepared figures and wrote the manuscript. F.S. established methods and the measurement strategy, performed experiments, analyzed data, prepared figures and the manuscript. N.E. and K.E. supervised the study, acquired funding, analyzed and interpreted data. N.A., P.K., C.C. and C.C. planned and performed the NTS experiments. O.G. and E.H. performed UNX-DOCA experiments. U.Z. contributed healthy human kidney tissue. C.D. and K.A. contributed archived human FFPE kidney biopsy material. All authors approved the final version of the manuscript.

